# PhenomNet: Bridging Phenotype-Genotype Gap: A CNN-LSTM Based Automatic Plant Root Anatomization System

**DOI:** 10.1101/2020.05.03.075184

**Authors:** Robail Yasrab, Michael P Pound, Andrew P French, Tony P Pridmore

## Abstract

This research will explore the phenotype-genotype gap by bringing two very diverse technologies together to predict plant characteristics. Currently, there are several studies and tools available for plant phenotype and genotype analysis. However, there is no existing single system that offers both capabilities in one package. Usually, Convolution Neural Networks used for plant phenotyping analysis and Recurrent Neural Networks used for genotype analysis. Both of these machine leanring methods require different input data for feature extraction, analysis and learning. Building a machine learning system for plant data that can make use of both graphic (for phenotype) and time-series (for genotype) is critical and challenging, especially when the system has to predict sensitive information regarding plant growth, accession and types. In this study, the proposed system will solve these problems by bringing two very different technologies, analysis methods and datasets. The proposed research aims to bridge the phenotype-genotype gap using CNN-LSTMs to process graphic and temporal data of plant roots. The proposed system “PhenomNet” offers segmentation of plant roots along with the classification of the given dataset into different accessions. The experiment results have shown that proposed CNN-LSTM architecture provides very high accuracy in comparison to manual or semi-automated approaches.

## I. Introduction

Plant phenotype and genotyping are terms associated with understanding plant growth and the effects of environmental conditions that influence the plant growth [1]. Plant phenotyping is an evolving biosciences area that helps to connect the plant genomics with agronomy and ecophysiology [2]. It is the study of biochemical and physical appears and characteristics of plants [3]. Whereas, plant genotyping is the study of plant age regression, mutation classification, genomic selection, disease resistance, etc. [4]. Several machine learning researchers worked plant phenotyping and genotyping separately, but there are very few who tried to put these two diverse but related paradigms together. Plant phenotype-genotype has improved dramatically due to the extensive use of new technology to accommodate the demand of food for the growing human population; there is a need for cultivating more advantageous genomic variants using precision breeding. This whole process depends heavily on a better comprehending of the phenotyping-genotyping relationship and filling the gap among these two key paradigms in plant science [4].

There are plenty of developments in bio-science, where image processing and machine learning-based tools used for manual or automatic plants feature extraction and analysis [5], [6], [7], [8], [9], [10]. Machine learning and computer vision played a vital role in the biosciences since the emergence of deep learning-based methods. In the past, many researchers [11], [9], [12], [13], [14] successfully applied computer vision-based methods for extracting, classifying and predicting plant characteristics and behaviors. Convolution Neural Networks (CNNs) [15] are one of the essential machine learning methods that usually applied for the classification and segmentation of plant image datasets [16], [17], [18]. CNNs are very good at the extraction of crucial features and classifying according to areas of interest. Therefore, CNNs are performing very well in terms of segmentation and classification of visual plant data (image, video). Plant genotyping analysis requires analysis of dynamic behavior and growth analysis [19]. CNNs are unable to model dynamic or temporal information (RNA/DNA sequences, textual data, etc.) accurately, so using static visual data to analyze genotype characteristics is not suitable enough. There is a need for additional temporal details to enrich the overall analysis. To resolve this issues, Recurrent Neural Networks (RNNs) [20] offers the capability to process time-series data. RNNs, especially Long Short-Term Memory (LSTM) [21] can process temporal data and very efficiently for dynamic analysis.

This study will combine CNN and LSTM to extract phenotyping and genotyping characteristics of plant roots. The proposed architecture will firstly use the CNN as an encoder to extract features to segment the visual attributes of given plant root images for phenotyping analysis. Later, these features forwarded to the LSTM decoder along with temporal information of the given root system. The LSTM decoder will process and output accurate classification and accession of the given root system. The Figure 1. shows a dual pipeline architecture, where Encoder-CNN will perform pixel-wise semantic segmentation of given root system and output complete root system architecture (RSA). The Decoder-LSTM take feature maps from final fully connected layers of the encoder and process these feature maps through hidden states to classify and predict the plant accession and age.

**Figure 1:**
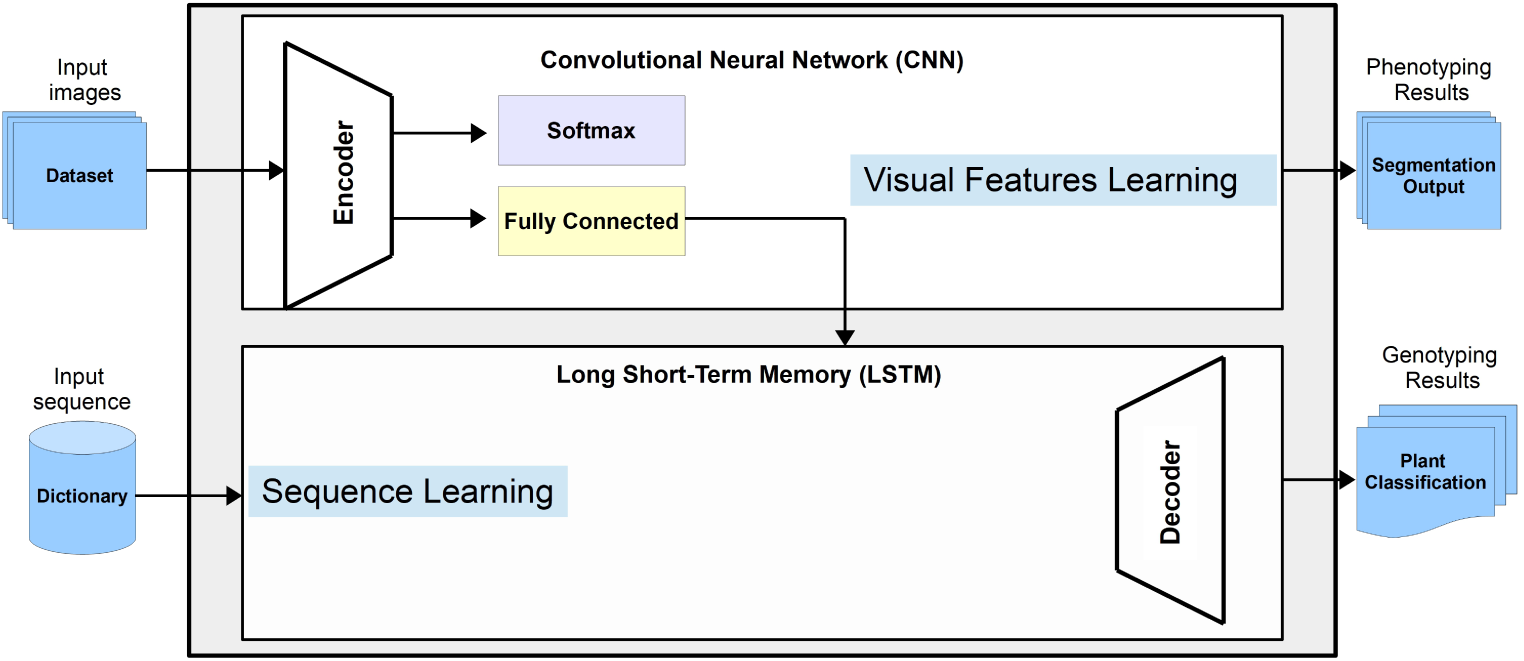
An overview of CNN-LSTM architecture. It is duel pipeline with encoder is a CNN that performs both segmentation of the root structure. The decoder (second pipeline) is LSTM that classify given room system into different treatments and accession groups.

The rest of the article outlined as follows: The next section describes the background of plant phenotyping and genotyping and its connection with machine learning. In the later section, we described the methodology of the proposed CNN-LSTM network and design. Later, the section based on different experiments performed to assess CNN-LSTM authenticity. The following part is about the results and analysis, and lastly, the conclusion and references section.

### A. Research Contributions

The main contribution of this research is that we have designed a state-of-the-art CNN-LSTM architecture that is capable of detecting and segmenting wheat roots. It can offer phenotyping and genotyping characteristics (a task often difficult for the traditional image-based phenotyping systems). Below is a summary of the key contributions of our research:

1. To the best of our knowledge, this is the first fully automated root phenotyping-genotyping system that allows complete root segmentation and classification without manual intervention.
2. Developed a novel dual-channel CNN-LSTM architecture that pixel-wise segments and classify the whole root system of wheat.
3. Benchmark results on publicly available wheat roots dataset.
4. Analyzed and compared the proposed CNN model with baseline semantic segmentation networks like VGG [22], FCN [23], SegNet [24], UNet [25], FastFCN [26].

## II. Background

An Artificial Neural Networks is a biologically inspired network with a fundamental computing unit, “Neuron.” Neuron basically takes multiple inputs, performs different computation and possibly outputs a non-linear function that is a weighted function of its input. A set of such artificial neural networks work together to build a computable artificial neural network system [27]. It composed of different layers and arithmetic functions. Fundamentally, a deep artificial neural network based on two or more layers. The first layer is called the input layer, the middle layer is called the hidden state/layer and the final layer is called the output layer. The way these layers connected determines the architecture of the model. The deep neural network architecture is trained for apparent traits to learn all the given features from the input. This training process involves making a forward pass of the input data several times, calculating the loss, updated each neuron weight, and finally making more and more accurate predictions [28].

Convolutional neural networks are composed of a trainable set of (aforementioned) layers. Each layer consists of a set of parameters; those can be trained from scratch or by a pre-trained CNN model (transfer learning [29]). CNN layers pass input to the next layer by processing it through convolution, non-linear activation (ReLU), pooling and normalization stages. The output of CNN is semantic class labels that represent the area of interest in the given input. CNNs specially designed to capture details from spatial data (such as images), and process this data through a set of layers (number varies from model to model) and finally produce predictions which could be either an object distribution (classification task), pixel-based classification (segmentation), or a single numeral value (regression problem). Since the introduction of CNNs, there are plenty of different varieties of CNN models that have emerged, ranging from shallow to ultra-deep CNN architectures, with a common goal to improve performance and accuracy for segmentation and classification tasks.

Recurrent neural network [20] is one of the popular types of Feed-Forward (FF) neural networks. In comparison to the standard FF neural network, RNN can deal with sequential input data. RNN uses internal memory to provide sequential/temporal information along with previously stored information for predictions. Standard RNN is unable to encode temporal dependencies that extend to more than a limited number of time steps [30]. LSTMs introduced by Schabber et al.[21] to resolve the abovementioned problem and capture time dependences in the given data. Along with resolving temporal dependencies, LSTM offers a solution for exploding/vanishing gradient by preserving and learning long time-based dependencies in data. LSTM is easier to manage, train and fine-tune as compared to CNNs. These can be effectively combined with CNN to offer a static-dynamic machine learning combination to resolve traditional classification problems using temporal data.

For the last few years, computer vision and machine learning-based systems provide a great deal of support for complex plant phenotyping tasks [11], [31], [32]. These systems help to extract useful features and information from plant data. These years computer vision-based methods are turning out to be essential for plant analysis. Plant phenotyping using machine learning identifies phenomic traits, measurements of quantitative parameters and physiological state of the plants [33]. The machine learning methods also used to assess genotyping aspects of plants like accession classification and growth analysis [19] — analysis of such dyanmic parameters related to plant physiology known as planted genotypes [34].

According to Siggh et al. [33], current machine learning methods used for identification, classification, quantification, and prediction of different phenomic tasks for plants. Deep learning and image processing are two methods effectively used for feature extraction from given plant images with higher accuracy and reliability. Most of the early approaches based on the rapid extraction of plant phenotypical information were related to the plant’s surface (leaf, steam, etc.). The proposed method attempts to apply machine vision to extract useful information about features of the below-ground plant (roots).

Processing temporal information along with traditional images based plant data to extract genomic and geometric information, is a very challenging task. CNNs work very well for classification and segmentation tasks, but they are architecturally not suitable for dynamic tasks due to their inability to encode the temporal dependencies. However, this problem can be addressed using RNN and specifically LSTMs, which is an improved version (of RNN) for handling dynamic inputs. Hence, using CNN along with LSTMs in plant phenotyping, becomes an emerging trend [35]. In this study, we proposed a CNN-LSTM architecture for plant classification for various accession groups. The framework can make use of temporal plant growth data to forecast the accession of a given plant. The proposed system specifically designed around a single plant with accession classification. Sakurai et al. [36] employ convolution LSTM to predict plant growth, though this study only limited to forecast plant age. In contrast, our proposed approach aimed to assess visual as well as accession features from the given plant image.

## III. Methodology

The proposed model intended to produce a phenotype-genotype description of the given input plant root images. The input to the proposed model is a set of images, ground truth and description of root system genotype characteristics. This model is unique as it aligns genotyping description to visual regions that described through multimodel embedding. These correspondences produced by CNN used as training data for a multimodel LSTM, which learns to generate genotype descriptions of given roots. Figure 4. offers a detailed insight into the proposed CNN-LSTM encoder-decoder architecture. This composed of a dual pipeline where encoder takes input image, processes it through the different layers and finally produces a semantic segmentation mask of root geometry. The decoder receives a data dictionary (holding genotype characteristics) as input along with encoder correspondences—this information processed through a given set of hidden states to produce root genotype characteristics finally.

**Figure 2:**
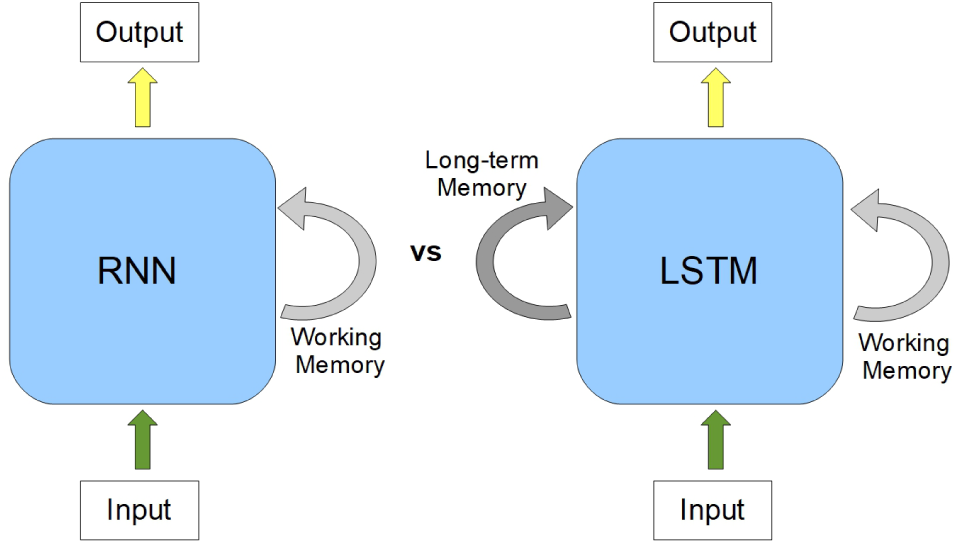
RNN v/s LSTM. a: RNNs use their internal state (memory) to process sequences of inputs, b: Long Short-Term Memory (LSTM) network is a varient of RNN, with addtional long term memory to remember past data.

**Figure 3:**
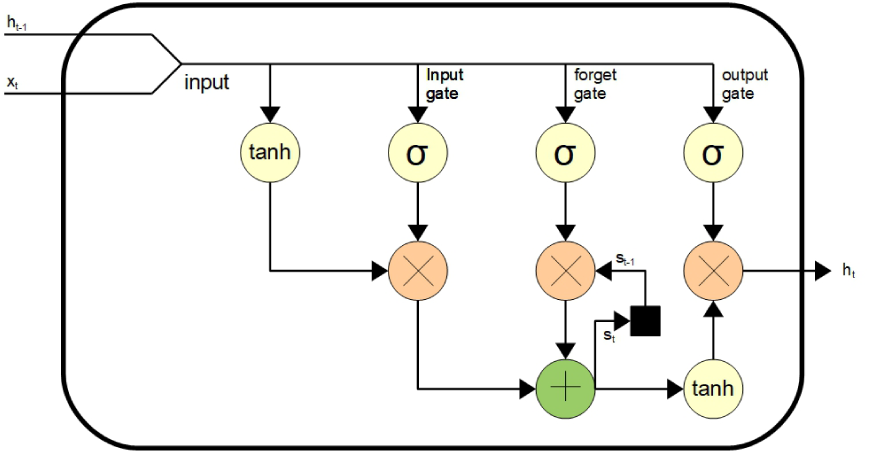
An unrolled LSTM. It takes input sequence *x*_*t*_ and *h*_*t*−1_ previous state, process this input through different gates (input, forget, out) and produce the output that could serve as final state or input for next hidden state.

**Figure 4:**
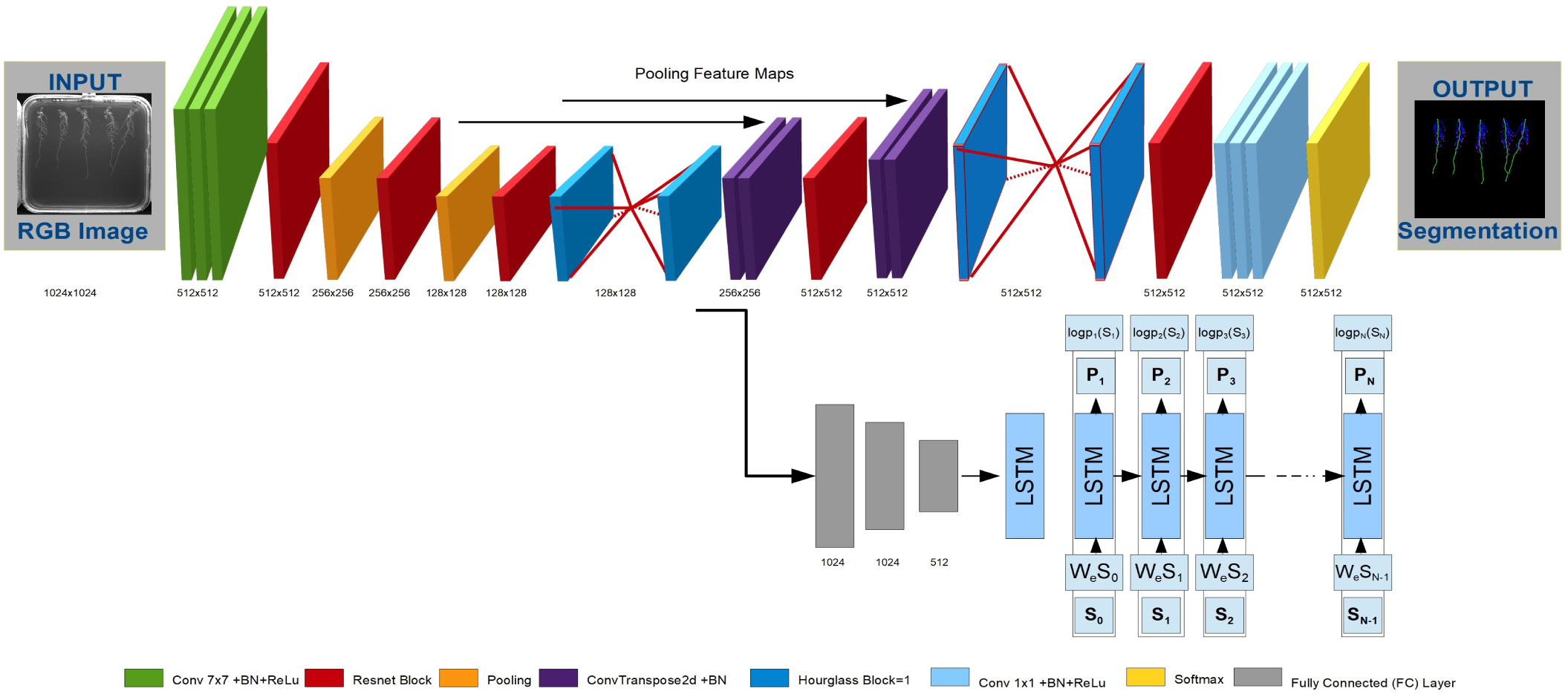
An overview of PhenomNet. An encoder-decoder network with encoder CNN and decoder LSTM. The encoder designed to pixel-wise segment given root image into a given set of classes. The decoder takes encoder correspondences and learns the genotype features of a given root system. The decoder outputs root genotype features like age and accession. Both networks work together to produce phenotype-genotype features at the same time. It is a simple network design that is end-to-end trainable.

### A. CNN-LSTM Network Architecture

#### 1) ENCODER CNN

To process sets of given input images of plant roots, we will use hourglass CNN [37] (with additional Residual (ResNet [38]) blocks and Fully Connected layers) as an encoder network. It is used to process given areas of interest (AoI) along with semantically segmenting phenotypical details. The encoder CNN produces root system architecture, including primary roots, lateral roots, seed location, root tips and background region (total six classes). The final layer of CNN encoder divided into two sub-pipeline. The first pipeline designed to produce semantic segmentation details of the given input image of the root: while the second pipeline adds additional activation and Fully Convolutional (FC) layers to produce sparse feature maps, which will be used as input for the LSTM-decoder.

#### 2) Decoder LSTM

The CNN encoder correspondences fed to LSTM decoder to predict genotype features as LSTMs can synthesize temporal dynamics in given image feature maps. The decoder LSTM classifies the plant roots through analyzing the sequences of features that are extracted from frames and by associated temporal variation. The primary purpose of using LSTM is to detect genotyping features of plant roots and capture the plant growth patterns over time. The genotype detector automatically classifies and predicts the plant age and accession by only giving plant images.

### B. Aligning genotyping and phenotyping data

The proposed CNN-RNN model is designed to produce genotyping-phenotyping information through a dual end-to-end trainable pipeline. This model inputs root images dataset to a CNN encoder to provide phenotyping information for given plant roots. Then, these CNN correspondences forwarded to decoder LSTM for learning genotyping features. Inspired by Karpathy et al. [39], the proposed model learns the ground dependencies tree relations to given root images with a ranking objective. Hence, the proposed models bind the genotyping descriptions to phenotype-based images regions in a standard multimodel embedding.

### C. DataSet

The primary dataset used in the proposed experiment is images of *Arabidopsis* (*A. thaliana*). This dataset is grown by Wilson et al. [40]. The dataset contains 4000 images of plates with each plate have five plants. Wells et al. [41] used near-infrared imaging methods to capture each plate image. The given dataset split into training and validation datasets with an 80/20 ratio.

#### 1) Growth Conditions

According to Wilson et al. [40] the seeds of Arabidopsis thaliana (L.) Heynh. (ecotype Columbia-0) were incubated in 5% (v/v) sodium hypochlorite for 5 min for the sake of surface-sterilized. Later, these seeds were washed three times with sterile water. Petri plates of size 125×125 used for sowing seeds in the vertical position. In this experiment, three key groups of Petri plates sown. The first group is planted with 60 ml 1/2 strength Murashige and Skoog media (Sigma) solidified with 1% (w/v) agar, group two with 1/4-MS media and group three with Full-MS media. Two days after sown at 4°C, the Petri plates transferred to controlled-environment chambers at 23C. This environment provides continuous light at a photon flux density of 150 mol m2 s1 for seven days. In this controlled environment, there are two categories of Petri plates with 12h and 24h photoperiod, each Petri plate marked with special treatment and photoperiod category for clear distinction from other plants [40]. A sample gentype distribution is shown in Figure 5 and a detailed dataset analysis is shown in Figure 6.

**Figure 5:**
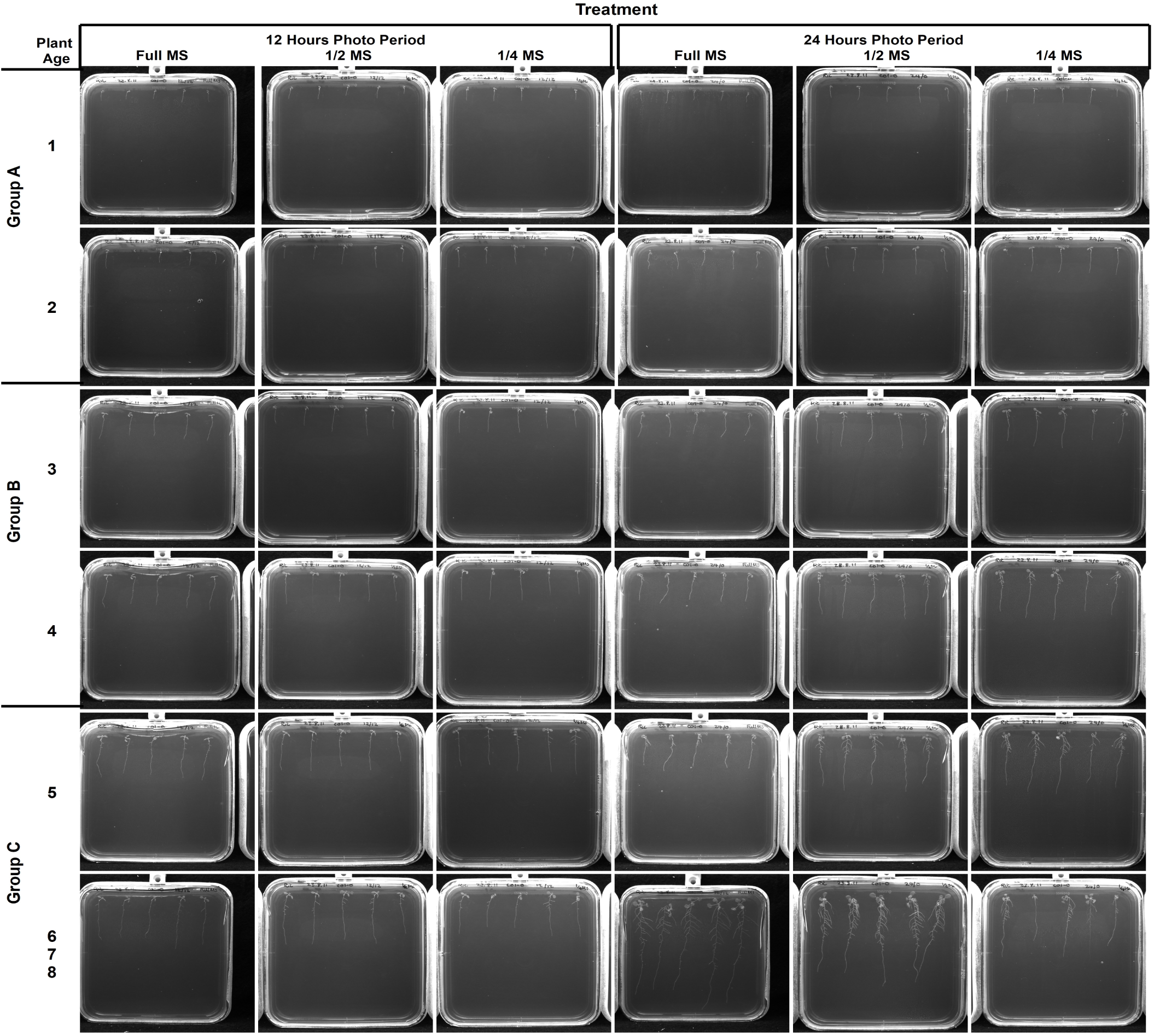
Example images from the dataset used during this work. *Arabidopsis* (*Arabidopsis thaliana*) seeds sown in Petri plates with three different (Full MS, Half MS, one forth of MS) strength of media (Murashige and Skoog media (Sigma) solidified with 1% (w/v) agar). The dataset divided into three accessions based on plant age. Plant age 1-2 week in Group-A, age 3-4 week in Group-B and age 5-8 weeks in Group-C.

**Figure 6:**
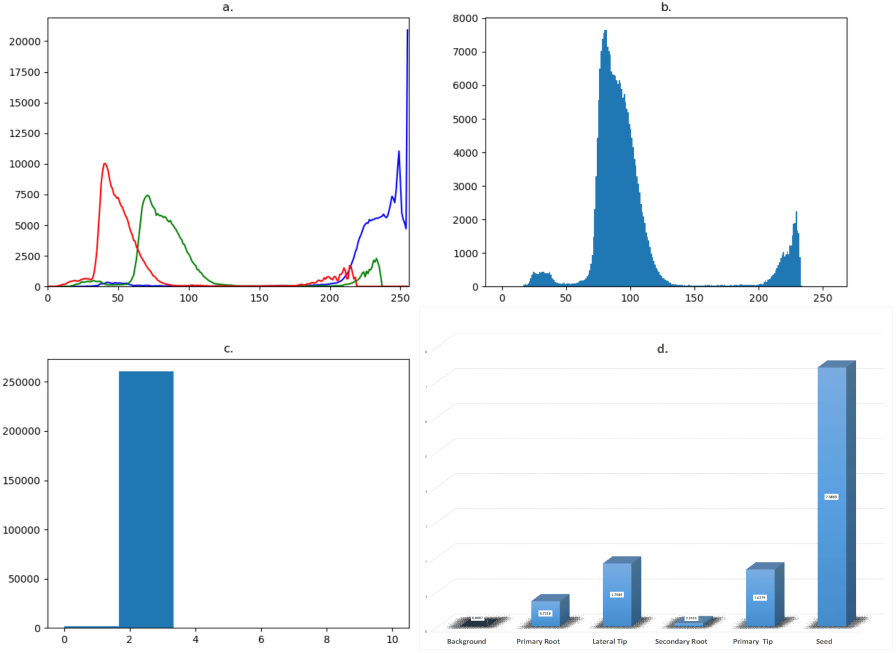
Input Dataset Analysis: a) The class distribution histogram of an RGB input image. It shows the histogram of red, blue and green colors. b) This graph shows a great deal of difference between the given set of foreground and background colors. Most of the areas covered in the input image is the gray background. c) It is a class distribution histogram of ground truth image. The figure is in the binary format, where background pixels are zero values, and foreground pixels are ones. d) An extensive class distribution difference shown in this class distribution histogram shows the class weights for each object of interest.

#### 2) Dataset Annotations

RootNav 2.0 [9] used to annotate the given plate images dataset; it is a fully-automated root phenotyping tool. It used to map and record the whole root system architecture (RSA). The annotated RSA is later saved in textual formate using Root System Markup Language (RSML) [42]. We have designed a special script to generate the ground truths (for CNN training) of given images using these RSML formate files.

### D. Dataset Dictionary

To record multiple genotype (temporal) features of each plant, we have created a dataset dictionary. This dataset dictionary holds the different temporal characteristics of each image in the given dataset. These characteristics include plant age, species, accession and specific treatments (as described in growth conditions section). The data dictionary is created manually with greater care to record all essential and correct details about each plant.

### E. Training CNN

The proposed CNN encoder trained to produce two very different types of feature maps for two very different purposes. The encoder acts as segmentation and classification architecture at the same time. The first pipeline (CNN) is intended to produce pixel-wise segmentation results for each input image. It is designed to segment primary roots, lateral roots, root tips, seed location and background. There are a total of six different segmentation classes that will be segmented by CNN architecture. The second pipeline (LSTM) acts as a classification network where it embedded with fully connected (FC) layers. This classification network intended to produce 512 feature maps that will be later forwarded to LSTM architecture to learn the temporal characteristics of plants. As LSTM work better with temporal sequences; therefore, these feature maps are feed to LSTM in the form of temporal sequences.

The size of the dataset plays a very significant role in the effective training and learning of CNN architecture. Bigger the dataset, better the training and prediction of a given set of classes. However, in our proposed experiment, we have a medium-sized dataset. In this scenario, we used some well-established machine learning proliferation methods to get effective results from a dataset. We have used transfer learning and data augmentation methods to improve the learning accuracy of the system.

#### 1) Transfer Learning

Transfer learning is a process of applying learned knowledge of one problem to a second (similar) problem. In deep learning, CNNs could be trained on a massive dataset with millions of images and later used for training some similar problems (dataset). It helps in learning a new task through transferring knowledge of related task that has already been learned [43]. In our case, we have used the pre-trained CNN model to transfer-learn a similar problem. The model used for transfer leanring was trained on a big dataset of Wheat (*Triticum aestivum* L.). The key reason to use this model is due to the apparent resemblance in roots structure. These dataset used for pre-training is composed of 3630 images of 1900×2000 pixels resolution. These images released by Pound et al. [17] and captured by Atkinson et al. [44]. The annotation and ground truth generation process is similar to the earlier dataset, where RootNav 1.0 was used to annotate areas of interest and save these mapping in RSML.

This RSML file used to generate ground truth for training and testing of the CNN system. The CNN is trained for 500 epochs until it starts converging to steady low loss and higher accuracy. The trained CNN model saved for further training of our similar dataset.

#### 2) Data Augmentation

Data augmentation is a method to adequately train CNN architecture when there is are not enough training images. In the proposed experiment, we used data augmentation to synthetically argument training dataset size. Horizontal-Fip and Random-Rotate are two well well-known argumentation methods were used in our experiments. The training images horizontally flipped with a 50% random rotate ratio. Images are also randomly rotated with a 15° degree rotation angle. The 15° degrees rotation angle set after several experiments; it is ensured that the original root system architecture not much influenced due to excessive rotation angel while learning process.

### F. LSTM Training

Decoder LSTM trained with encoder generated feature maps to learn temporal characteristics. The CNN encoder produces 512 output feature maps through final FC layers that are transferred to LSTM to predict temporal information about every given input image. The decoder LSTM is designed with additional (deeper) neurons to offer better learning. This provides an enhanced capability to LSTM for learning complicated temporal details about the input dataset and also helps to enhance accuracy and overall learning process.

## IV. Experiment and Results

### A. Training Setup

We have used PyTorch [45] deep learning library for implementation and Nvidia GPU with two 12GB memory used for training. Due to the high-resolution size of the input and extensive CNN-LSTM parameters, we cannot go beyond batch size one on a single GPU. Cross-Entropy Loss with Adam optimization function used for loss and accuracy optimization. The initial learning rate was set to 0.1 and decreased by a factor of 10 at every 100 iterations. The whole architecture trained for 500 epochs until the loss and accuracy of the system start converging and stay steady after 500 epochs. The system further trained for an additional 50 epochs to make sure a stable convergence.

### B. Weighted Loss

The proposed CNN and LSTM systems produce two very different outputs. The decoder LSTM architecture uses cross-entropy loss for producing genotype features of the given plants. The CNN architecture with a softmax final layer to produced pixel-wise segmentation results using cross cross-entropy loss with a weighted-loss function to facilitate CNN architecture for better training [46]. Plant roots have a complex and cluttered nature, and there is also a visible unbalanced between the number of foregrounds and background pixels. Loss calculated on the unbalanced dataset can cause a bias toward background pixel and leads to lower accuracy in the segmentation process. The class balancing approach with the standard cross-entropy loss function helps us for better training and quick converging to a given set of classes. Weights assigned to each class inversely proportional to the medium frequency in which that class appears throughout in entire training dataset. It means a class that appears more often will have lower weight (e.g., background class), and class that appears less frequently will have higher rates (e.g., seed class).

### C. Extraction of Phenotype/Genotype Characteristic

The proposed LSTM encoder-decoder architecture intended to segment and classify a given plate-root system. The CNN encoder equipped with a final softmax layer that segment the root system into the primary root, lateral root, root tips and seed categories. The second pipeline of the network feeds feature maps from encoder CNN to fully connected layer and later to the LSTM architecture. This LSTM network learns given temporal details of the root system. Finally, it classifies the root system for different phenotype features like accession, age and treatment. The given dataset is composed of a sequence of plate images grown over days. These plates categorized based on different accession and treatment. Each plate has received different treatments for growth analysis. Therefore, we have listed all possible classes of the accession and treatment parameter into the dataset directory. Developing this data directory is a very critical task that needs to be done with additional care for the accurate classification of plants. The given plants divided into three accession and three treatment classes. There are a total of three different (accession and treatment) genotype classification of the dataset. This section will present the results and analysis of the proposed experiment. It will outline different CNN and LSTM architectures that analyzed for getting the best performance for pixel-wise segmentation and classification of temporal features. The results assessed in terms of accuracy and loss factors. This analysis conducted among different CNN/LSTM trained architectures. The intended systems trained with the same dataset and hypermeters (number of epochs, training loss, batch size, etc.). These are different CNN architectures used for experiments; for example, VGG [22], FCN [23], SegNet [24], UNet [25], and FastFCN [26] and proposed Pheno-Net. These CNN networks compared in terms of the number of parameters, accuracy and loss. The LSTM trained with different neurons depths to check the most suitable depth for the learning of the given dataset dictionary. These models implemented for parallel processing using PyTorch Machine Learning Library and Python.

The Pheno-Net architecture has a massive number of trainable parameters; it composed of CNN and LSTM joined in a single end-to-end trainable pipeline. So, it adds millions of trainable parameters that lead to extensive training and GPU memory consumption. In this scenario, to train this architecture with high-resolution images on a single GPU (12Gb size) is not possible. For this reason, there is a need for CNN architecture with more memory efficiency, fewer parameters and higher trainable accuracy. During analysis, it realized that proposed Pheno-Net architecture performs well not only in terms of memory but also in terms of higher accuracy with reduced loss. The proposed PhenoNet architecture is composed of multiple convolution layers, pooling layers, residual blocks, an hourglass module and final softmax layer. It offers a better performance in comparison to other benchmark CNNs architectures in conjunction with chosen LSTM architecture.

To assess the most appropriate LSTM arrangement for experiments, we trained LSTMs with different neuron depths (hidden states). It started with 128 neuron depth (number of neurons for LSTM’s hidden state) and kept moving-up till 512 neuron depth. After evaluating the final results, it realized that LSTMs with 512 neuron depth offered us much better learning capabilities in comparison to less deep neurons LSTMs. Therefore 512 hidden neuron layers finalized for dual pipeline architecture of the proposed system. We combined these architectures and trained the given dataset through a single end-to-end trainable design for 500 epochs.

This architecture trained until the loss and accuracy start converging and hold steady learning status. Statistical analysis performed among semi-automated annotations and manual directory building parameters. It assessed that the proposed network results are almost similar in terms of accuracy and lower loss in comparison to the benchmark architecture.

Table I shows the comparison of different CNN architectures. In this scenario, we have compared VGG [22], FCN [23], SegNet [24], UNet [25], and FastFCN [26] along with our proposed PhenoNet. The results have shown that the suggested design is smaller in terms of other architectures and offers almost similar/higher accuracy and lower loss with better performance. It provides very economical trainable parameters and improves performance in terms of enhancing the learning capabilities of the network.

**Table I:** Quantitative Comparison: This graph shows a quantitative comparison of propsoed CNN with well-know earlier CNN architectures for roots segmentation problem. propsoed CNN outperforms all the other systems. We have used global average, class average and mIoU as standard metrics for comparison of segmentation results.

| CNN Arch. | Class avg. | Global avg. | mIoU |
| --- | --- | --- | --- |
| VGG [22] | 35.47 | 41.27 | 29.04 |
| FCN [23] | 39.24 | 46.55 | 37.32 |
| SegNet [24] | 61.74 | 78.44 | 46.76 |
| UNet [25] | 74.14 | 79.00 | 50.16 |
| FastFCN [26] | 82.00 | 81.35 | 56.22 |
| **Proposed CNN** | **90.10** | **97.00** | **64.33** |

The Table II presents the comparison between different LSTM depth architectures. We experimented with LSTM neuron depth 128 to 512 LSTM depth. The given results showed that LSTM with depth 512 offers the most improved outcomes and better capability in terms of classifying the temporal characteristics of given plants. It also noticed that in comparison with other CNN architectures, the LSTMs depth 512 offers the most better results.

**Table II:** Quantitative Comparison of different LSTM Models: this table shows a quantitative comparison of different LSTM models trained to learn the given genotype features of plant roots. It proved that deep hidden state LSTM performs exceptionally well as compared to shallow models.

| Model A LSTM | Model B LSTM | Model C LSTM |
| --- | --- | --- |
| 1 Hidden Layer | 2 Hidden Layer | 3 Hidden Layer |
| 512 Hidden Units | 256 Hidden Units | 128 Hidden Units |
| **96.05%** | **95.24%** | **91.22%** |

The Figure 7 shows genotype-phenotype results from the experiments. It shows the color GT (segmentation) and classification outcomes that describe the given input images in a much more detailed way. This Figure 8 presents the loss and accuracy curve for the training and testing stages of the proposed architecture. This curve represents the loss of the proposed architecture goes down as the training epochs go up and it stays steady after 600k iterations. The proposed architecture train well in terms of improvement in accuracy and performance.

**Figure 7:**
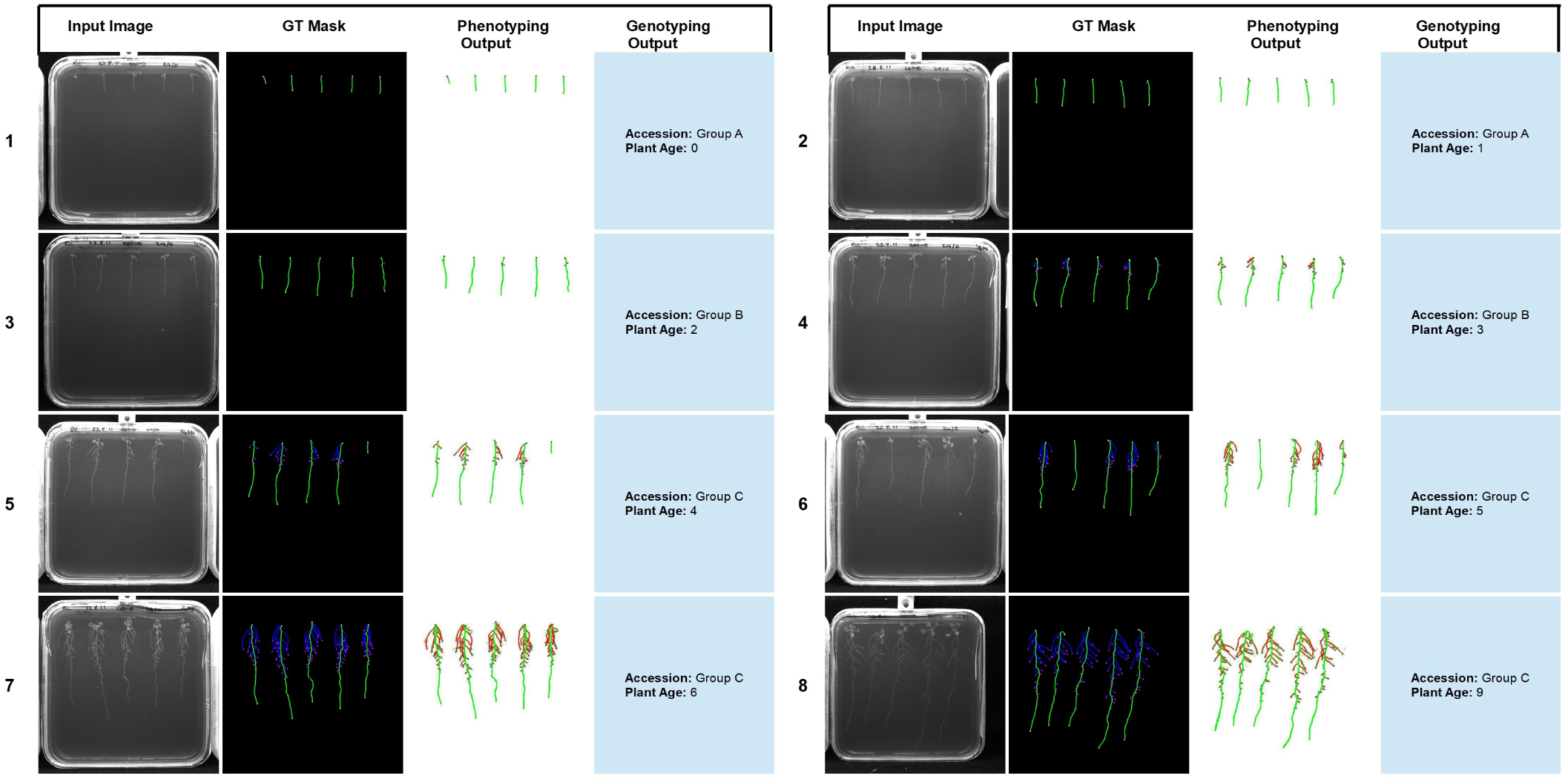
Example outputs from *PhenomNet*. (a) Input image. (b) Colour-coded GT mask. (c) Color-Coded CNN output segmentation masks. (d) LSTM classification output.

**Figure 8:**
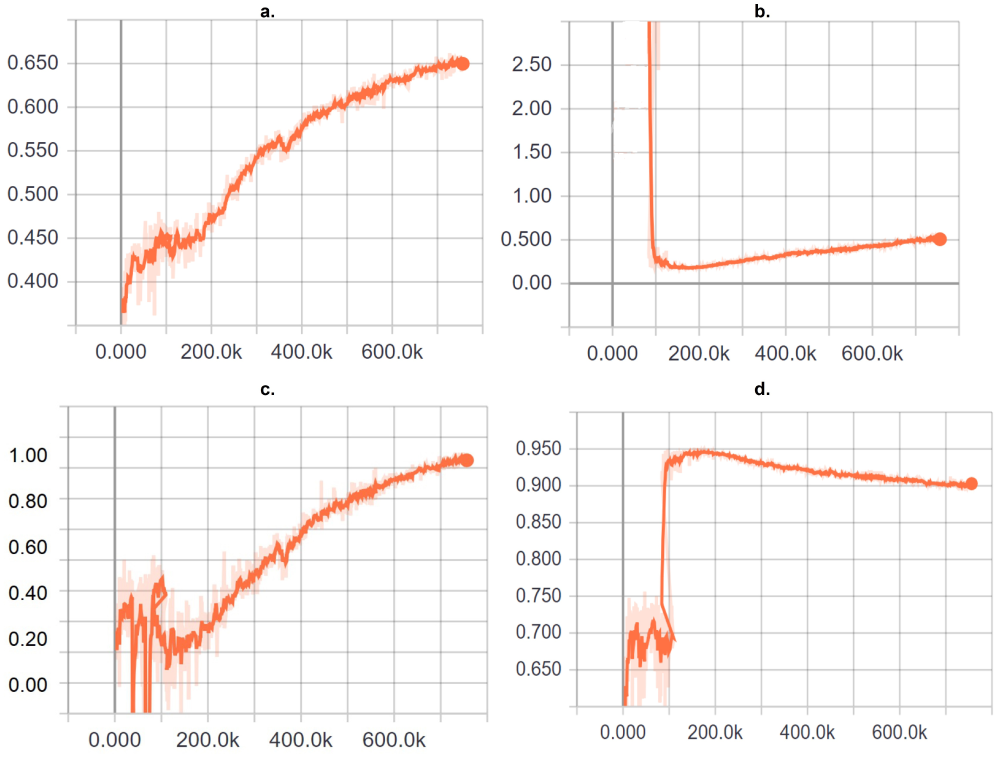
Graphical Analysis of Results: These graphs show a quantitative comparison of CNN training and validation results. The training curves are in the background, while validation curves are highlighted. a). This graph shows the Mean IoU of the proposed system. This highest value attained by the proposed system during the training and validation process. b). This graph shows the training and validation Loss of the proposed method. c). Overall Accuracy of training and Validation datasets. d) Mean Accuracy of training and Validation datasets.

### D. Discussion

The idea of this research was to train dual pipeline end-to-end trainable CNN-LSTM architecture that is capable of processing images and temporal sequences at the same time. The proposed research results have proved that we can segment and classy plant root system architecture at the same time with higher accuracy. The basic idea is to make use of the power of machine learning systems to present and forecast several dimensions in plant phenotyping. This trained model can be used for further analysis of similar images. It can also be extended to other datasets with a considerable amount of training. The code and trained models will be available online on GitHub for open-source analysis and usage. In the future, we aimed to extend the overall capability to the proposed system to further (new) dataset of different plant species. Another key feature we aimed to add is to save the RSA to Root System Markup Language (RSML) for better analysis of root systems after classification and segmentation.

## V. Conclusion

This paper has presented the idea to bridge the gap between plant phenotype and genotype. The aim is to design an architecture that could be trained end-to-end and it can automatically classify and extract the genotype-phenotype features at the same time. So the proposed architecture is an automated framework designed to extract and segment plant root features. It also learns temporal cues and predicts the growth pattern as well as accession information about plant roots. The proposed framework is an encoded decoder architecture whose encoder based on the convolution neural network, while the decoder based on an LSTM. The encode CNN acts as a segmentation block. In contrast, the decoder LSTM acts as a classification setup to predict the ascension growth of given plant roots. To the best of our knowledge, it is the first kind of architecture that attempted to bridge the gap between genetic and phenotyping characteristics of plant roots.

It has shown that the proposed architecture is a fully automatic architecture that performs almost equivalent to a semi-automate (RootNav1) or handcrafted features extraction methods for plant genotyping and phenotype. For training purposes, we have used different datasets (transfer learning process) to enhance the learning capability of the proposed system. The final results proved that the proposed architecture could process any complicated and challenging root systems and forecast their geometry, ascension and growth patterns excellently. Furthermore, the proposed architecture could be extended for further genotyping analysis like disease detections or environmental conditions analysis. Another key feature of the proposed system is an end to end single pipeline architecture that can be trained on any given dataset. In the future, we aimed to extend the current application of CNN-LSTM to further datasets. We also aimed to train LSTM with more comprehensive genotype characteristics.

